# A tale of winglets: evolution of flight morphology in stick insects

**DOI:** 10.1101/774067

**Authors:** Yu Zeng, Conner O’Malley, Sonal Singhal, Faszly Rahim, Sehoon Park, Xin Chen, Robert Dudley

**Affiliations:** Department of Integrative Biology, University of California, Berkeley, CA 92870, USA; Schmid College of Science and Technology, Chapman University, Orange, CA 92866, USA; Department of Biology, CSU Dominguez Hills, Carson, CA 90747 USA; Islamic Science Institute, Universiti Sains Islam Malaysia, 71800 Bandar Baru Nilai, Negeri Sembilan, Malaysia; Centre for Insect Systematics, Universiti Kebangsaan Malaysia, 43600 Bangi, Selangor, Malaysia; Department of Biology, The College of Staten Island, The City University of New York, NY 10314, USA; Department of Biology, The Graduate School and University Center, The City University of New York, NY 10016, USA; Smithsonian Tropical Research Institute, Balboa, Republic of Panama

**Keywords:** body size, evolution, flight, phasmid, sexual dimorphism, wing size

## Abstract

The evolutionary transition between winglessness and a full-winged morphology requires selective advantage for intermediate forms. Conversely, repeated secondary wing reductions among the pterygotes indicates relaxation of such selection. However, evolutionary trajectories of such transitions are not well characterized. The stick insects (Phasmatodea) exhibit diverse wing sizes at both interspecific and intersexual levels, and thus provide a system for examining how selection on flight capability, along with other selective forces, drives the evolution of flight-related morphology. Here, we examine variation in relevant morphology for stick insects using data from 1100+ individuals representing 765 species. Although wing size varies along a continuous spectrum, taxa with either long or miniaturized wings are the most common, whereas those with intermediate-sized wings are relatively rare. In a morphological space defined by wing and body size, the aerodynamically relevant parameter termed wing loading (the average pressure exerted on the air by the wings) varies according to sex-specific scaling laws; volant but also flightless forms are the most common outcomes in both sexes. Using phylogenetically-informed analyses, we show that relative wing size and body size are inversely correlated in long-winged insects regardless of sexual differences in morphology and ecology. These results demonstrate the diversity of flight-related morphology in stick insects, and also provide a general framework for addressing evolutionary coupling between wing and body dimensions. We also find indirect evidence for a ‘fitness valley’ associated with intermediate-sized wings, suggesting relatively rapid evolutionary transitions between wingless and volant forms.

## 1. Introduction

Flight is fundamental to the ecology and evolutionary diversification of pterygote insects by allowing for three-dimensional mobility and greater access to nutritional resources (Dudley, 2000). Nonetheless, approximately 5% of the extant pterygote fauna is flightless (Roff, 1994), and various conditions of reduced wing size (e.g., brachyptery and microptery) are found across the neopteran orders. Given structural costs and high energy expenditure during flight, maintenance of the flight apparatus is not universally favored by selection. Partial reduction or complete loss of wings is associated with various morphological and ecological factors, such as developmental tradeoffs, enhanced female fecundity, and reduced demand for aerial mobility in certain habitats (Roff, 1990; Roff, 1994). In these cases, smaller wings exhibit reduced aerodynamic capability, but may serve secondarily derived non-aerodynamic functions such as use in protection, stridulation, and startle displays (see Dudley, 2000).

Wing evolution can also be influenced indirectly by selection on overall body size. Generally, reduced body mass enables greater maneuverability in flight (e.g., more rapid translational and rotational accelerations), although numerous factors influence insect size evolution (see Blanckenhorn, 2000; Chown and Gaston, 2010). Furthermore, both flight capacity and body size can be subject to sex-specific selection. As a consequence, sexual size dimorphism (SSD) is typically associated with intersexual niche divergence and with sexual selection (see Shine, 1989; Hedrick and Temeles, 1989). Sexual wing dimorphism (SWD) can in some cases be decoupled from SSD, and may be associated with divergence in aerial niche and wing use (e.g., DeVries et al., 2010). Selection for greater locomotor capacity in males can lead to male-biased SWD, and also to female-biased SSD (see Roff, 1986). It is therefore of interest to consider patterns of sexual dimorphism in both wing and body size within a phylogenetic context.

The stick insects (Phasmatodea) exhibit great diversity in both wing and body size (**Figs. 1, 2**), but underlying evolutionary patterns are not well characterized. Most winged stick insects possess rudimentary and tegmenized forewings. Phasmid hindwings (designated ‘wings’ hereafter) can be of various sizes and exhibit expanded cubital and anal venation with well-developed flight membranes. Fossil evidence suggest that both wing pairs were full-sized in ancestral stick insects (see Shang et al., 2011; Wang et al., 2014; Yang et al., 2019), whereas numerous extant species exhibit wing reduction. Earlier studies have proposed frequent evolutionary transitions between winged and wingless morphologies, although the directionality and the detailed dynamics of phasmid wing evolution remain contested (see Whiting et. al., 2003; Stone and French, 2003; Trueman et al., 2004; Whiting and Whiting, 2004; Goldberg and Igić, 2008). Nevertheless, size-reduced wings must lead to degradation in aerodynamic performance, with possibly concurrent changes in body length and mass. Given the unresolved history of wing size evolution of this group, we use the term ‘reduction’ to describe wings that are developmentally truncated relative to a full-sized morphology, without assessing the directionality of wing size evolution within the phylogeny.

**Figure 1.**
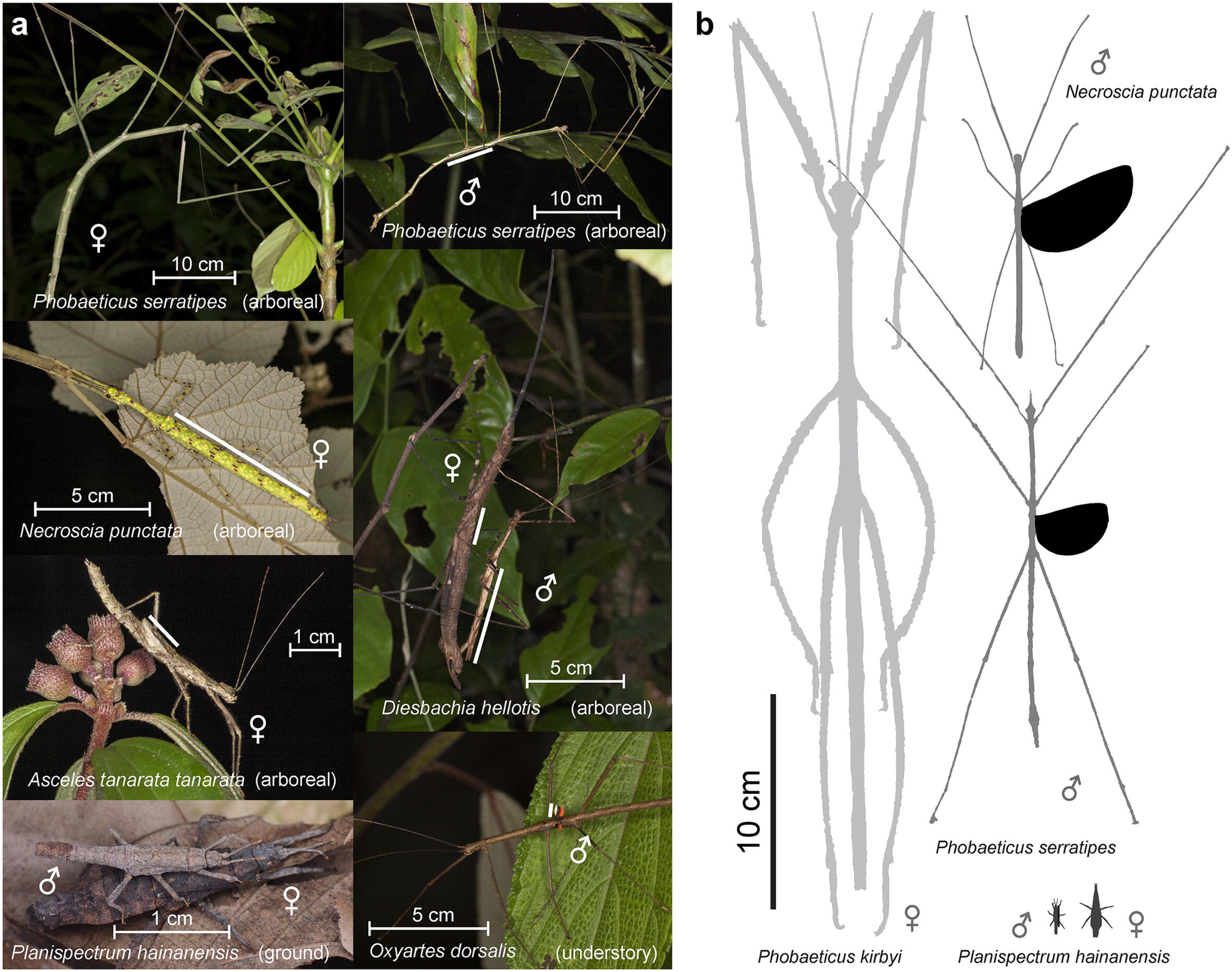
Diversity of flight morphology in stick insects as shown by variation in wing and body size. (a) Examples of stick insects with different sized wings; white line segments indicate hindwing length in winged taxa. (b) Spectrum of interspecific variation in body and relative wing size for representative species. (Photo of *Planispectrum hainanensis* courtesy of Chao Wu.)

**Figure 2.**
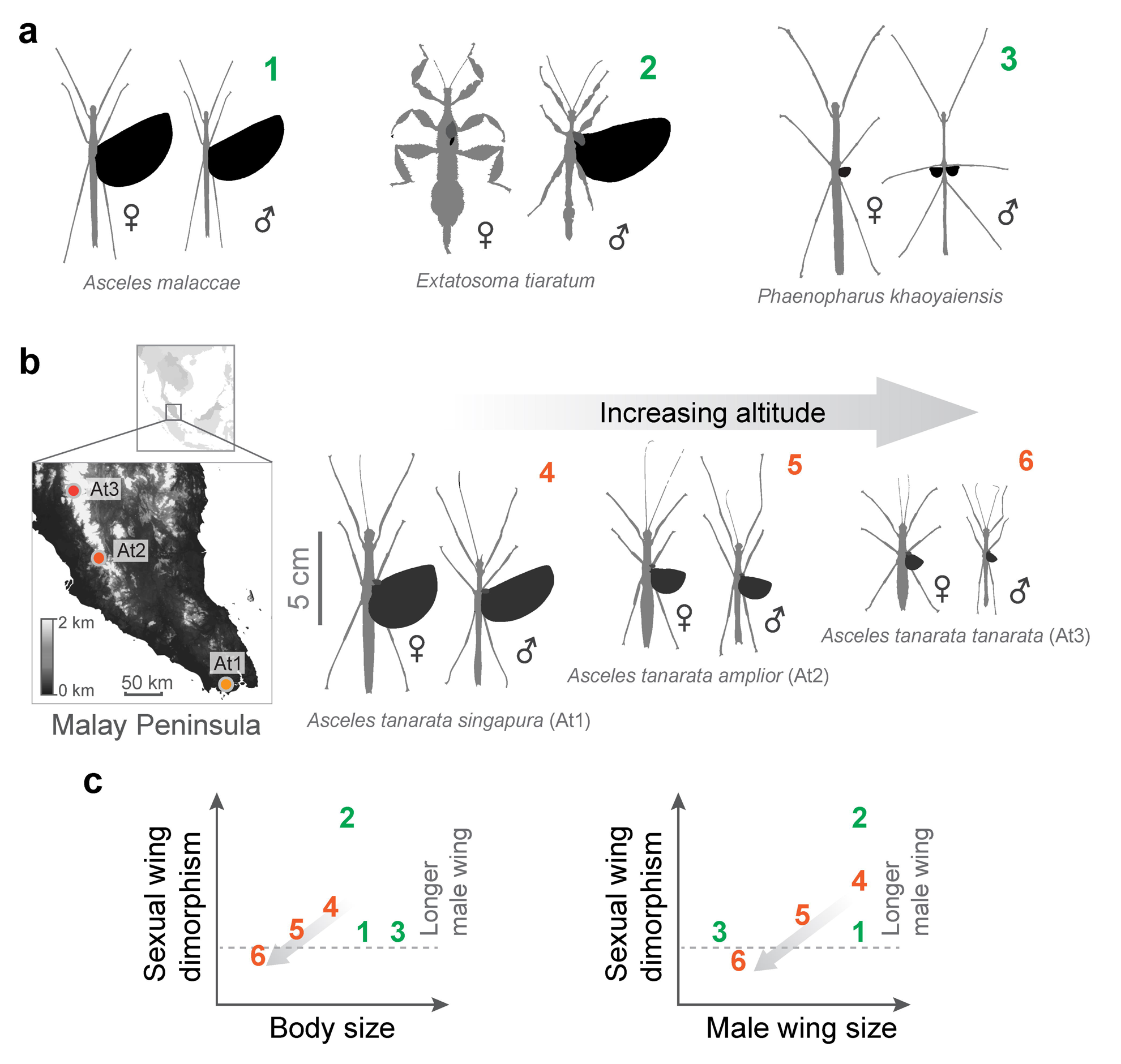
Sexual wing dimorphism (SWD) in stick insects. (a) Representative combinations of variable relative wing size and SWD: (1) low SWD with long wings in both sexes, (2) extreme SWD with long wings in males only, and (3) low SWD with short wings in both sexes. (b) Variation in wing and body sizes for the *Asceles tanarata* species group, for which SWD transitions from male- to female-biased with increasing altitude. (c) Schematic demonstration of variations in SWD with respect to body size and male wing size; numbers denoting taxa depicted in (a) and (b). The gray arrow indicates elevational changes with increasing altitude in the *A. tanarata* group.

Here, we examine the evolution of phasmid flight morphology on a macroevolutionary scale. We <u>first</u> describe variation in wing and body size using data from 1100+ individuals across 765 species, including intraspecific data from the *Asceles tanarata* species group with three subspecies exhibiting altitudinal variation in both wing and body size (see Brock, 1999; Seow-Choen and Brock, 1999; **Fig. 2b**). This group represents one of the few well-documented cases of features of insect flight morphology being distinctly correlated with a gradient in environmental parameters. Second, using live specimens, we derive a scaling relationship for wing loading with wing and body size. We then use this empirical model to extrapolate variation in flight capability across all sampled species. Third, we use phylogenetic correlational analyses to quantify the relationship between changes in wing size (reflecting flight ability) and body size through evolutionary time. Throughout these analyses, we also assess overall patterns of sexual dimorphism among phasmid species within phylogenetic and allometric contexts. In particular, we examine the correlation between SWD and SSD to address the role of sex-specific ecology in driving the diversity of flight morphology. For example, if selection on male-biased mobility and on female-biased fecundity were coupled, we might expect an inverse correlation between SWD and SSD.

## 2. Materials and Methods

### Morphometrics

Our sampling primarily focused on winged phasmid clades, given available data (see **Supplementary Fig. S1**). Wing length (L_w_) and body length (L) data were primarily obtained from literature sources, and were enriched with measurements on both captive-reared and field-collected insects (see section ‘Scaling of wing loading’). Taxonomic justifications followed Phasmida Species File (Brock, 2019), as downloaded and formatted using custom-written scripts in MatLab (Supplementary Materials). For the *A. tanarata* species group, male and females of three subspecies were collected in the field (see also Brock, 1999). The main dataset includes measurements on 599 males and 533 females from 765 species (∼23% of 3348 known species), of which 367 species included data on both sexes (**Supplementary Dataset 1**). If available, mean measurements were used; otherwise, median values were calculated based on ranges between maximum and minimum values. The relative wing size (Q) was defined as the ratio of wing length to body length: 

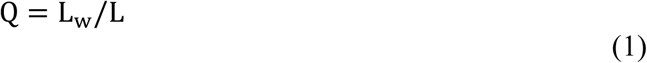

To avoid ambiguity, hereafter the term ‘wing size’ primarily refers to the relative wing size. SWD (ΔQ) was calculated as: 

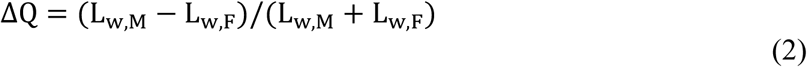

where the subscripts M and F denote male and female, respectively. The sign and magnitude of ΔQ thus represent the type and level of SWD. For example, ΔQ < 0 represents female-biased SWD, ΔQ = 0 represents absence of SWD, and ΔQ = 1 when the female is wingless and the male is winged. Similarly, SSD (ΔL) was calculated as: 

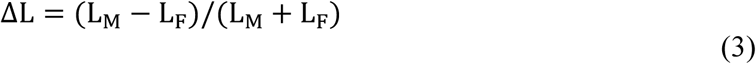

### Scaling of wing loading

The loss of aerodynamic capability was assessed using the parameter wing loading, the ratio of body weight to total wing area. First, the scaling of body mass relative to body length can be expressed as: 

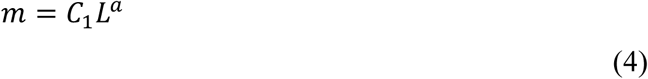

where *C*_*1*_ is the slope coefficient and *a* is the scaling exponent. If body mass varies under isometry (i.e., body geometry is conserved across different body sizes), we expect *a* to equal 3. Similarly, the power-law scaling of total wing area (A_w_) with Q can be expressed as: 

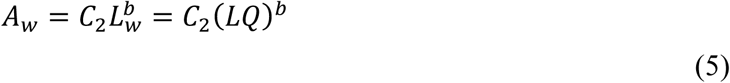

where *C*_*2*_ is the slope coefficient and *b* is the scaling exponent. If wing area varies under isometry, we expect *b* to equal 2. Combining Eqns. 4 and 5, we have the power-law scaling of wing loading (p_w_) with L and Q: 

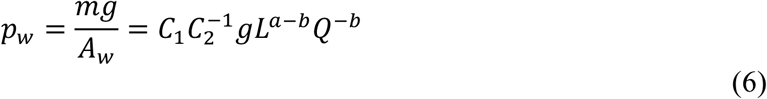

We then developed empirical models of the scaling of wing loading for both sexes based on Eqns. 4-6. We sampled total wing area (A_w_), body mass (m) and wing length (L_w_) from 23 males and 21 females of field-collected and captive-bred insects from 36 phasmid species (**Supplementary Dataset 2**). Digital images were obtained dorsally for insects placed on horizontal surfaces with all legs laterally extended; projected areas of fully unfolded wings were manually extracted using Photoshop (CS6, Adobe Systems Inc., San Jose, CA, USA). Values for A_w_, L_w_ and L were obtained using ImageJ (Schneider et al., 2012).

### Bayesian phylogenetic reconstruction

We used three mitochondrial genes (cytochrome oxidase subunit I (COI) gene, cytochrome oxidase subunit II (COII) gene, and large subunit rRNA (28S) gene; total length 2149 bp) and one nuclear gene (histone subunit 3 (H3) gene; 350 bp) (primer details in **Supplementary Table S1**). Our molecular sequencing covered nine species, including all three taxa from the *A. tanarata* group (**Supplementary Dataset 3**). We extracted total genomic DNA from leg tissue using a modified high-salt protocol (Aljanabi and Martinez, 1997) and, subsequently, quantified and diluted the DNA using a Nanodrop spectrometer. We amplified each loci using standard PCR conditions. Amplified products were cleaned with Exosap and sequenced using PCR primers with BigDye v3.1 on an Applied Biosystems 3730 machine. For other species, we downloaded sequence data from the same four genes from GenBank. Our molecular data set covers about 70% of the recognized tribes of Phasmatodea (Brock et al., 2019) and two outgroup species (Embioptera), with 95% of the species > 95% complete by locus.

Sequences were assembled in Geneious (v6.1.7, Biomatters) and aligned using the MUSCLE algorithm (Edgar, 2004). Gene alignments were checked manually for accuracy. jModelTest v0.1.1 was used to determine the best fitting substitution model for each gene based on the Akaike Information Criterion (AIC) (Posada, 2008). Next, we estimated a time-calibrated phylogeny in BEAST package (v1.10.4; Drummond et al., 2012). Across genes, we used unlinked substitution models and linked clock and tree models. To date the phylogeny, we used the fossil crown group phasmid *Renphasma sinica* dated 122 Myr ago (Nel and Delfosse, 2011) to set the minimum age of the divergence between Embioptera and Phasmatodea. Also, we included two fossil calibrations, following Buckley et al. (2008). Fossil Euphasmatodean eggs from mid-Cretaceous dated to 95–110 Myr ago were used (see Rasnitsyn and Ross, 2000; Grimaldi and Engel, 2005) to determine the age of the most recent ancestor of Euphasmatodea. The sister group relationship between *Timema* and Euphasmatodea has been confirmed by both morphological and molecular evidence (Whiting et al., 2003; Bradler, 2009). Therefore, we assumed the divergence between Euphasmatodea and *Timema* occurred more than 95 Myr ago. Furthermore, we used fossil leaf insect dated 47 Myr ago (Wedmann et al., 2007) and fossil eggs of Anisomorphini dated 44 Myr ago (Sellick, 1994) to set the minimum age of the nodes of the most recent common ancestors of leaf insects and Pseudophasmatinae, respectively. We first optimized the Markov chain Monte Carlo (MCMC) operator by performing short runs (1 × 10^7^ cycles) with a relaxed lognormal model and a Yule model, and adjusted the operators as suggested by the program. Then, we ran ten analyses for 2 × 10^8^ generations each. We monitored convergence and determined the burn-in using TRACER v1.7 (Rambaut et al., 2018). After discarding burn-in (25%), we used a maximum credibility approach to infer the consensus tree in TreeAnnotator v1.10.4.

### Phylogenetic correlations

A total of five morphological traits was used in phylogenetic analyses (L and Q of both sexes and sex-averaged L, ΔQ, and ΔL). First, we calculated the phylogenetic signals (λ) for all characters using the maximum-likelihood approach implemented in Phytools (Pagel, 1999; Revell, 2012). This model was compared with alternative models where *λ* was forced to be 1 or 0 in order to find the best-fitting model. The best-fitting model was found using the likelihood ratio (LR) test, 

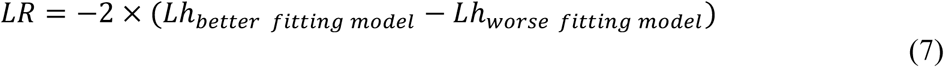

whereby the better fitting model has the highest log-likelihood score, *Lh* (Pagel, 1997, 1999; Freckleton et al., 2002). When *λ* = 0, this suggests trait evolution is independent of phylogenetic association, which is equivalent to generalized least square (GLS) model. We also assessed the evolutionary distribution of morphological traits with maximum-likelihood ancestral state reconstruction, using ‘fastAnc’ function in Phytools (Revell, 2012).

For the species that lacked molecular data, we added them as polytomous tips to the node representing the latest common ancestor on the tree. We then generated 100 random trees with randomly resolved polytomous tips. Each new node was added using the function ‘multi2di’ (package ‘ape’; Paradis et al. 2004), and was given a branch length that was randomly drawn from a normal distribution of branch lengths with a mean of 0.1 × mean branch lengths of the original tree, and a standard deviation of 0.01 × the standard deviation of branch lengths from the original tree.

We analyzed phylogenetically-justified correlations using phylogenetic generalized least square (PGLS) analyses (package ‘caper’; Orme et al., 2013). For each correlation, we ran PGLS on all random trees and summarized the results (ML*λ* and coefficients), which were then compared with those from ordinary generalized least square (GLS) tests conducted without referring to the phylogeny (i.e., *λ* = 0). To avoid zero-inflation in correlational analyses due to winglessness (i.e., Q = 0), we used two methods for correlations involving Q: (1) excluding species with Q = 0; and (2) converting Q to a pseudo-continuous ordinal variable as: 1 (Q = 0), 2 (0 < Q < 0.3), 3 (0.3 < Q < 0.6), or 4 (Q > 0.6; see Symonds and Blomberg, 2014). Also, we adopted a similar protocol for all correlations involving ΔQ, whereby ΔQ was converted to: 1 (ΔQ < 0), 2 (ΔQ = 0), 3 (0 < ΔQ < 0.3), 4 (0.3 < ΔQ < 0.6), or 5 (ΔQ > 0.6). In addition, to accommodate the bimodal distribution of Q (see Results), we categorized short-winged and long-winged insect groups as ‘0’ and ‘1’, and applied logistic regression models separately. We defined short- and long-winged characters using the distribution of Q values across all species (see dotted line in **Fig. 3b**).

**Figure 3.**
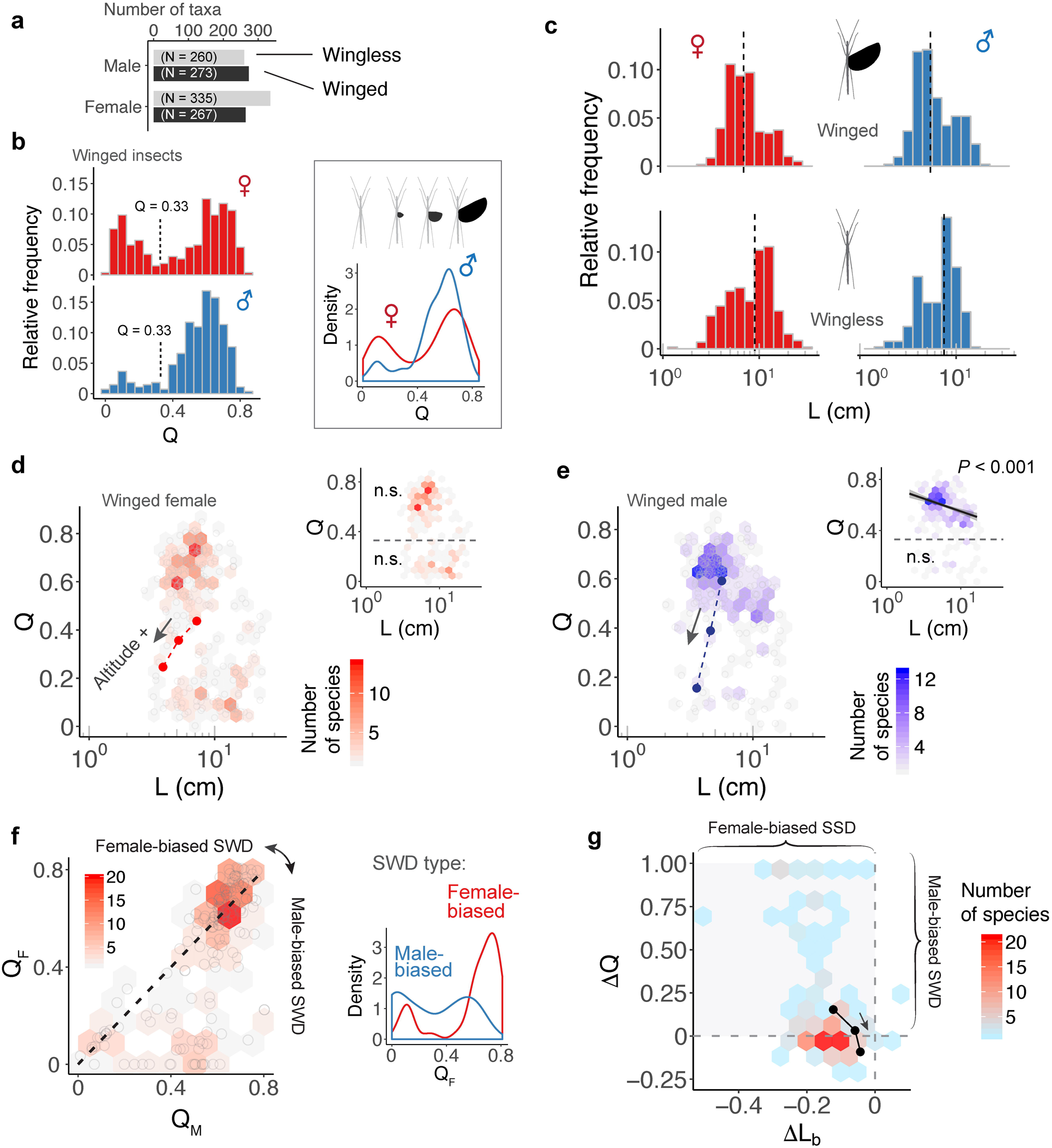
Variations in wing and body size among stick insect species. (a) Number of winged and wingless species, as grouped by the two sexes. (b) Relative frequency distribution and density of relative wing size (Q) for winged taxa. The vertical dashed lines indicate a region of phenotypic space that few species occupy. (c) Relative frequency distribution of body size (L). Black dashed lines denote median values. (d) – (e) Scatter plots of Q versus L for winged stick insects, indicating bimodal distributions. The color of overlaid hexagons represents the number of species, as scaled by the heat map inset. Insets show results of generalized least squares (GLS) regression models (trend line with 95% C.I.) for short- and long-winged insects, as categorized by the cutoff Q defined in (a). An inverse correlation between Q and L is found in long-winged males. (f) Scatter plot of female relative wing size versus male relative wing size (*N* = 183 species). The color of overlaid hexagons represents the number of species for each parameter combination. The majority (80%) of female-biased SWD (i.e., the area above the dashed line) is associated with medium-length to long wings (Q_F_ > 0.5), as indicated by the increasing density of female-biased SWD in long-winged females (Q > 0.4) (inset). (g) Scatter plot of SWD index versus SSD index, showing the predominance of female-biased SSD and continuous variation in SWD (57% male-biased and 43% female-biased). For (d), (e), and (g), the three dark dots represent subspecies of the *A. tanarata* group, showing sex-specific trends of wing and body size reduction with increasing altitude (as indicated by arrows).

## 3. Results

### Sex-specific variation in flight-related morphology

Among all sampled insects, ∼44% of females and ∼51% of males were winged. Relative wing size (Q) varied continuously from complete winglessness (Q = 0) to fully-sized wings (i.e., Q ≈ 0.85; **Fig. 3a,b**). For both sexes, the relative frequency of Q was bimodally distributed with a valley near Q = 0.3, and two peaks near Q = 0.1 and Q = 0.7, respectively. Variation in the bimodal distribution was sex-specific, whereby the majority of males exhibited medium- to fully-sized wings (i.e., Q > 0.4) whereas most females exhibited either medium- to fully-sized wings, or alternatively miniaturized wings (Q < 0.3). The frequency distribution of body length (L) was bell-shaped, with females exhibiting a wider range and greater mean length compared to males (male range: 1.7 cm – 19.0 cm, female range: 1.3 cm – 28.5 cm; male mean: 6.92 cm, female mean: 8.71 cm; see **Fig. 3c**). In both sexes, the median body length of the wingless group was greater than that of the winged group. The GLS regression model suggested a significant inverse correlation between Q and L in long-winged males, but not in other groups (**Fig. 3d,e**).

At the species level, extent of SWD varied with relative wing size. Of 183 winged species with data from both sexes, ∼57% (88 species) showed various levels of male-biased SWD (ΔQ > 0; **Fig. 3f**). Female-biased SWD, however, tended to be found in species for which both sexes possessed long wings. Of the other 42% of species with female-biased SWD (ΔQ < 0), most exhibited long wings (Q > 0.6 in both sexes). In general, phasmids showed different combinations of a continuously varying SWD and female-biased SSD (**Fig. 3g**). For *A. tanarata* group, the reduction in coefficients of wing and body size toward higher altitudes was sex-specific (**Fig. 3d,e**). Males showed a relatively higher extent of wing reduction, leading to a reversal of SWD from male- to female-biased (**Fig. 3g**).

### Sex-specific flight reduction

Scaling of wing area with wing length was nearly isometric, with an exponent (*b*) of approximately 1.84 in both sexes (**Fig. 4a**; **Table 1**). The allometric scaling of insect mass with respect to body length was, however, sex-dependent, with females exhibiting a higher slope coefficient relative to males (**Fig. 4b**). Larger female phasmids thus have disproportionally greater mass. Consequentially, the allometric coefficient for wing loading in females (i.e., 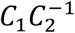) was ∼1.5 greater than that of males (Eqn. 6; **Fig. 4c,d**). Females generally have much greater wing loading and potentially greater loss of aerodynamic capability when compared to males of the same relative wing size. This may partially underlie the high frequency of female-biased SWD found in long-winged taxa (see Discussion). For males, variation of body size plays little role in the variation of wing loading. With the scaling exponent for body size (i.e., *a* − *b*; see Eqn. 6) approximately equal to zero, wing loading in males is exclusively dependent on relative wing size (Q). Notably, the male of *Heteropteryx dilatata*, a morphological outlier with full-sized forewings, exhibits a higher wing loading relative to other males due to its disproportionally greater body mass.

**Table 1.** Comparison of coefficients for the allometric scaling of body mass (m) and the scaling of wing area (A_w_) with wing length (L_w_) (see Eqns. 4 and 5). Values represent means with 1 s.e. in brackets. Units are consistent with those in text and figures: L, *cm*; m, *g*; L_w_, *cm*; A_w_, *cm*^*2*^.

| Sex | Models and parameters |  |  |  |
| --- | --- | --- | --- | --- |
| | $m = C_1 L^a$ | | $A_w = C_2 L_w^b$ | |
| | $C_1$ | $a$ | $C_2$ | $b$ |
| **F** | -2.09 (0.19) | 2.38 (0.22) | -0.16 (0.04) | 1.84 (0.07) |
| **M** | -2.08 (0.14) | 1.84 (0.16) | -0.24 (0.03) | 1.84 (0.06) |

**Figure 4.**
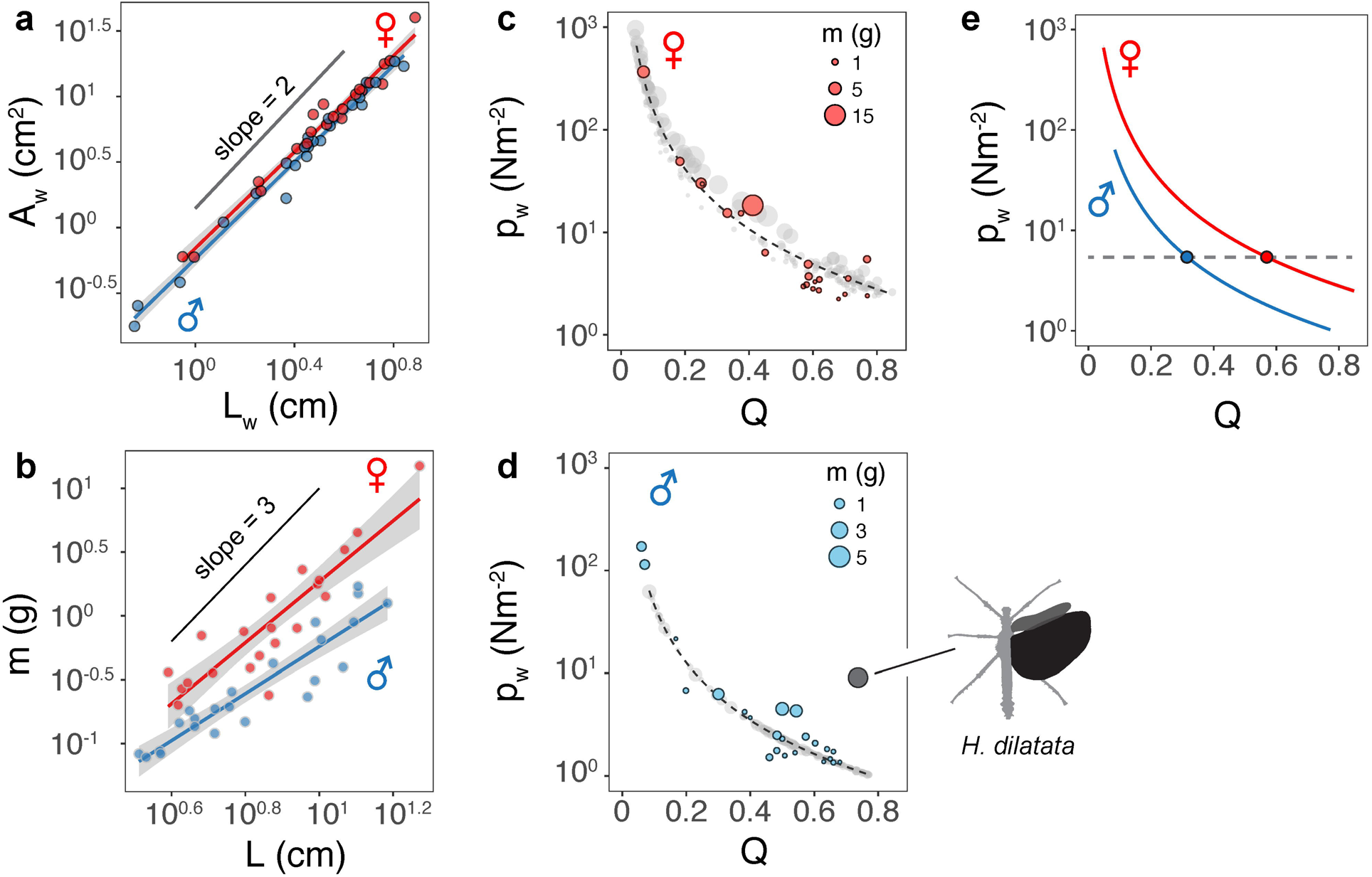
Scaling of flight-related morphology. (a) Near-isometric scaling of wing area with relative wing size. Trend lines are based on linear regression models with slopes equal to 1.84±0.05 and 1.84±0.07 (mean±s.e.m.) for males and females, respectively; R^2^ = 0.98 and *P* < 0.0001 for both sex groups). (b) Allometric scaling of insect body mass with body length. Trend lines are based on linear regression models; Males: slope = 1.84±0.16, R^2^ = 0.86, *P* < 0.0001, females: slope = 2.39±0.21, R^2^ = 0.86, *P* < 0.0001. (c) – (d) Allometric scaling of wing loading (p_w_) in females and males, respectively. Colored dots represent insects for which body mass and wing area were directly measured. Gray dots are estimates based on wing and body lengths using regression models (see Methods). Trendlines are based on a logistic fit. The regression model for males omitted *Heteropteryx dilatata* (dark gray dot), which is a morphological outlier with well-developed forewings. (e) Comparison of the scaling of p_w_ with respect to Q between two sexes, showing that disproportionally longer wings in females are required to attain wing loading equivalent to that of males.

Variation in wing loading can also be presented as a three-dimensional landscape relative to wing and body size. The allometric effect is stronger in females, whereas males exhibit a smaller lower boundary for wing loading without any allometric effect (**Fig. 5a,b**). Projecting the species richness distribution onto these landscapes demonstrates a clustering of taxa on the wing loading functional landscape (**Fig. 5c,d**). Both sexes showed two major clusters associated with low and high wing loadings, corresponding to long-winged and miniaturized-wing morphologies, respectively. The majority of long-winged females were allometrically constrained to values of wing loading between 10^−0.5^ Nm^-2^ < p_w_ < 1 Nm^-2^, whereas long-winged males clustered near a value of 10^−1^ Nm^-2^, with a number of taxa characterized by even lower values. The miniaturized-wing taxa in both sexes tended to concentrate within the high wing loading regime (i.e., p_w_ > 10 Nm^-2^). Despite sexual differences in the topology of the wing loading landscape, a threshold wing loading between 1 Nm^-2^ < p_w_ < 10 Nm^-2^ was associated with a largely unoccupied region of phenotypic space (i.e., Q = 0.3; **Fig. 3b**).

**Figure 5.**
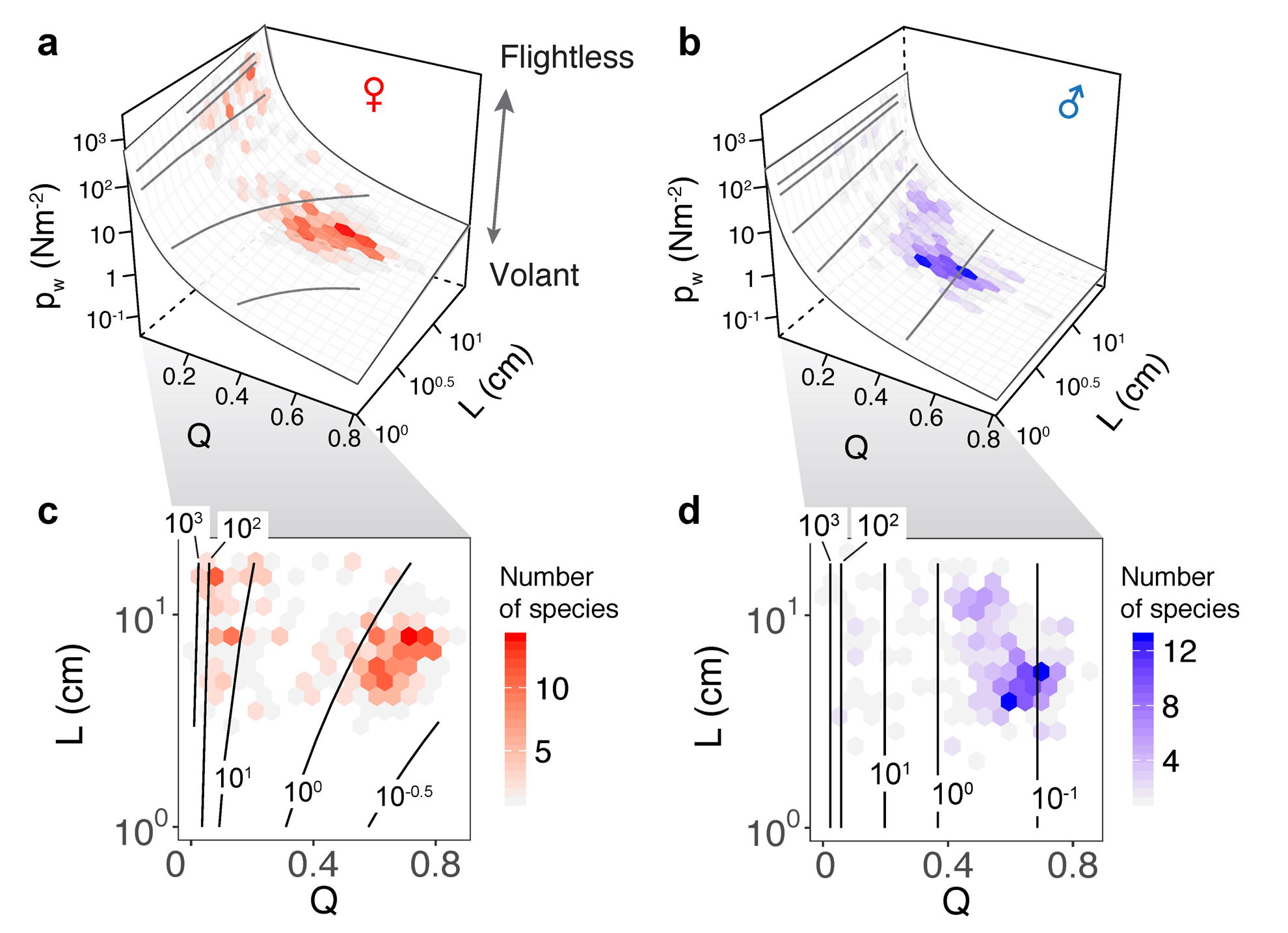
Sex-specific landscapes of wing loading relative to dimorphism and body length among phasmid species. **Fig. 5a** and **Fig. 5b**, wing loading relationships for females and males, respectively between the two sexes; females typically have higher wing loading than males and stronger allometric effects relative to body length. **Fig. 5c** and **Fig. 5d**, contours of the wing loading landscape for females and males, respectively, as overlaid with hexagonal bins for species counts (**Fig. 3d**); wing loading distribution differs substantially between the sexes.

### Wing size-dependent evolutionary correlations

Our tree topology and estimates of diversification times were largely concordant with those of published phasmid phylogenies (see Whiting et al. 2003; Bradler et al. 2015; Robertson et al. 2018; **Fig. 6; Supplementary Fig. S2**). Within the *A. tanarata* group, the divergence time between the lowland subspecies (*A. tanarata singapura*) and two highland subspecies was ∼ 3 Myr ago, whereas the divergence time between two highland subspecies was ∼1 Myr ago. Significant phylogenetic signal was present in all morphological traits (**Table 2**). Our conservative ancestral state reconstruction showed high evolutionary lability of wing and body size, and suggested that an intermediate relative wing size (Q < 0.4) preceded various levels of gains and losses in both sexes (**Supplementary Fig. S4**).

**Table 2.** Summary of statistical results for best model fits, comparing phylogenetic generalized least squares (PGLS) models (λ estimated by maximum likelihood, ML) with generalized least square (GLS) models (λ = 0) for log-transformed body length (i.e., log_10_ L) and relative wing size (Q). Species-wise traits were analyzed for all taxa using available data for both sexes.

| | Variable | $N$ | $ML\lambda$ | $Lh$ (PGLS) | $Lh$ (GLS) | $Lh$ (PIC) | PGLS vs. GLS | |
| --- | --- | --- | --- | --- | --- | --- | --- | --- |
| | | | | | | | $LR$ | $P$ |
| **Male** | $Q_M$ | 533 | 1.001 | 293.3 | -111.9 | 227.5 | -810.3 | < 0.0001 |
| | $\log_{10}(L_M)$ | 533 | 0.922 | 317.9 | 108.9 | -91.4 | -418 | < 0.0001 |
| **Female** | $Q_F$ | 597 | 1.001 | 401.1 | -100.4 | 292.7 | -1002.9 | < 0.0001 |
| | $\log_{10}(L_F)$ | 597 | 0.916 | 271.8 | 76.1 | -140.1 | -391.4 | < 0.0001 |
| **Species-wise comparison** | Sex-average $L$ | 367 | 0.967 | -1701.9 | -1846.7 | -1920.5 | -289.5 | < 0.0001 |
| | $\Delta L$ | 367 | 0.576 | 442.9 | 431.4 | 88.6 | -23 | < 0.0001 |
| | Sex-average $Q$ | 367 | 0.949 | 114.5 | -59.1 | 137.2 | -347.2 | < 0.0001 |
| | $\Delta Q$ | 367 | 0.926 | 12.6 | -60.2 | -3.5 | -145.6 | < 0.0001 |

**Figure 6.**
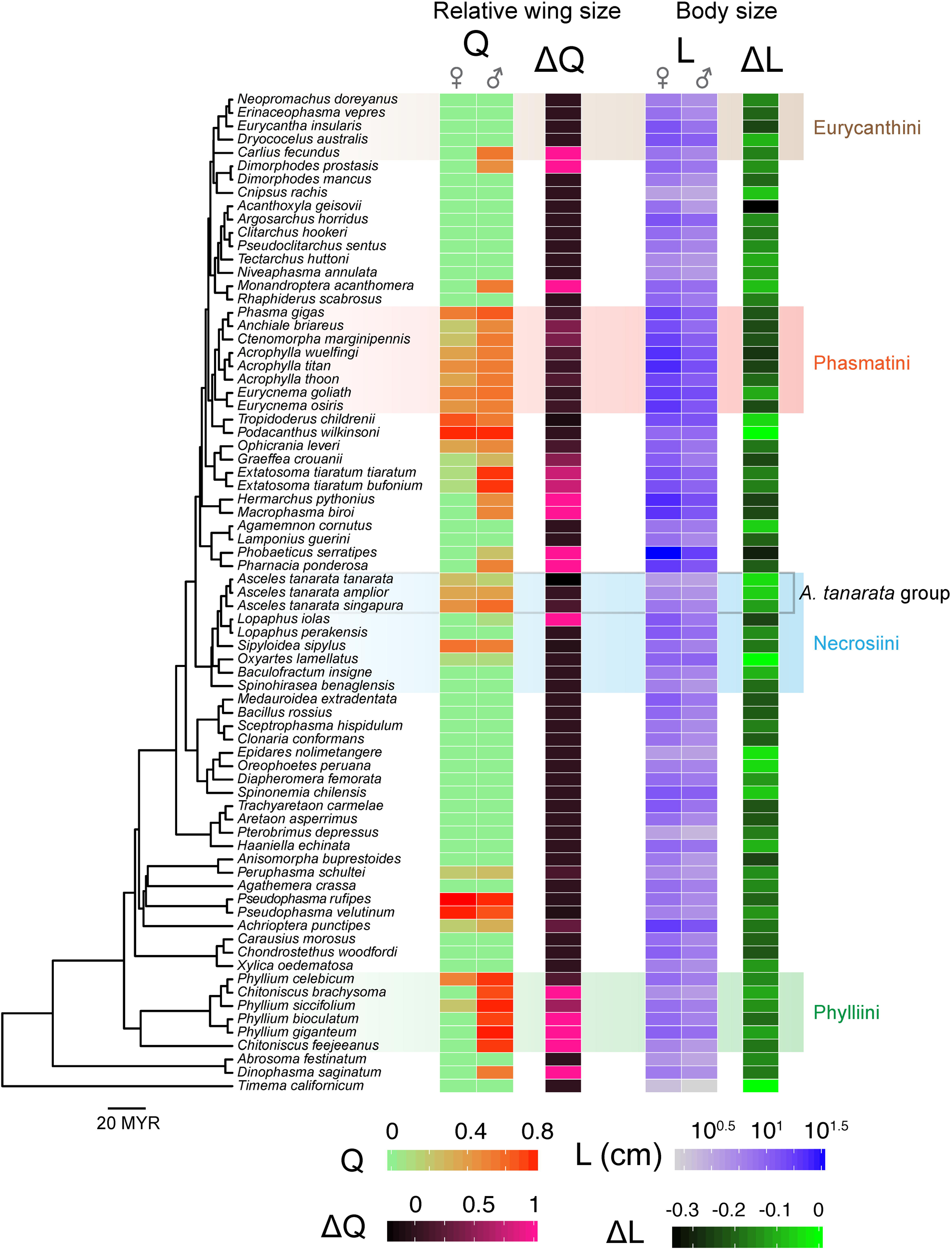
Phylogenetic relationships among sampled taxa, with flight-related morphology annotated on tree tips. Tree topology is based on concatenated COI, COII, H3 and 28S data (see Methods), and branch lengths are proportional to time since divergence (in millions of years). The tree is pruned to show a selection of species with data obtained for both sexes; selected tribes are highlighted to show variation within groups (see **Supplementary Fig. S2** for the complete tree).

Based on the PGLS results, there was a significant inverse correlation between Q and L in long-winged insects (Q > 0.33) of both sexes (**Fig. 7a,b; Table 3**), which supported our initial hypothesis on evolutionary coupling between wing and body size. In addition, sex-averaged body size was coupled with the extent of both SWD and SSD in long-winged species (Q > 0.33 in both sexes), suggesting opposite trends of variation in SWD and SSD along the gradient of sex-averaged body size (**Fig. 7c**). An exemplar of this correlation is demonstrated in **Fig. 7d,e**, whereby increases in both SSD and SWD lead to greater sexual differences in wing loading. Short-winged insects generally lacked significant correlations between wing and body size (**Fig. 7a,c**). Across all winged species, variation in female traits contributed substantially more to intersexual differences, as shown by the predominant roles of female Q and L values in determining variation in SWD and SSD, respectively (**Supplementary Fig. S6, Table S2**).

**Table 3.**
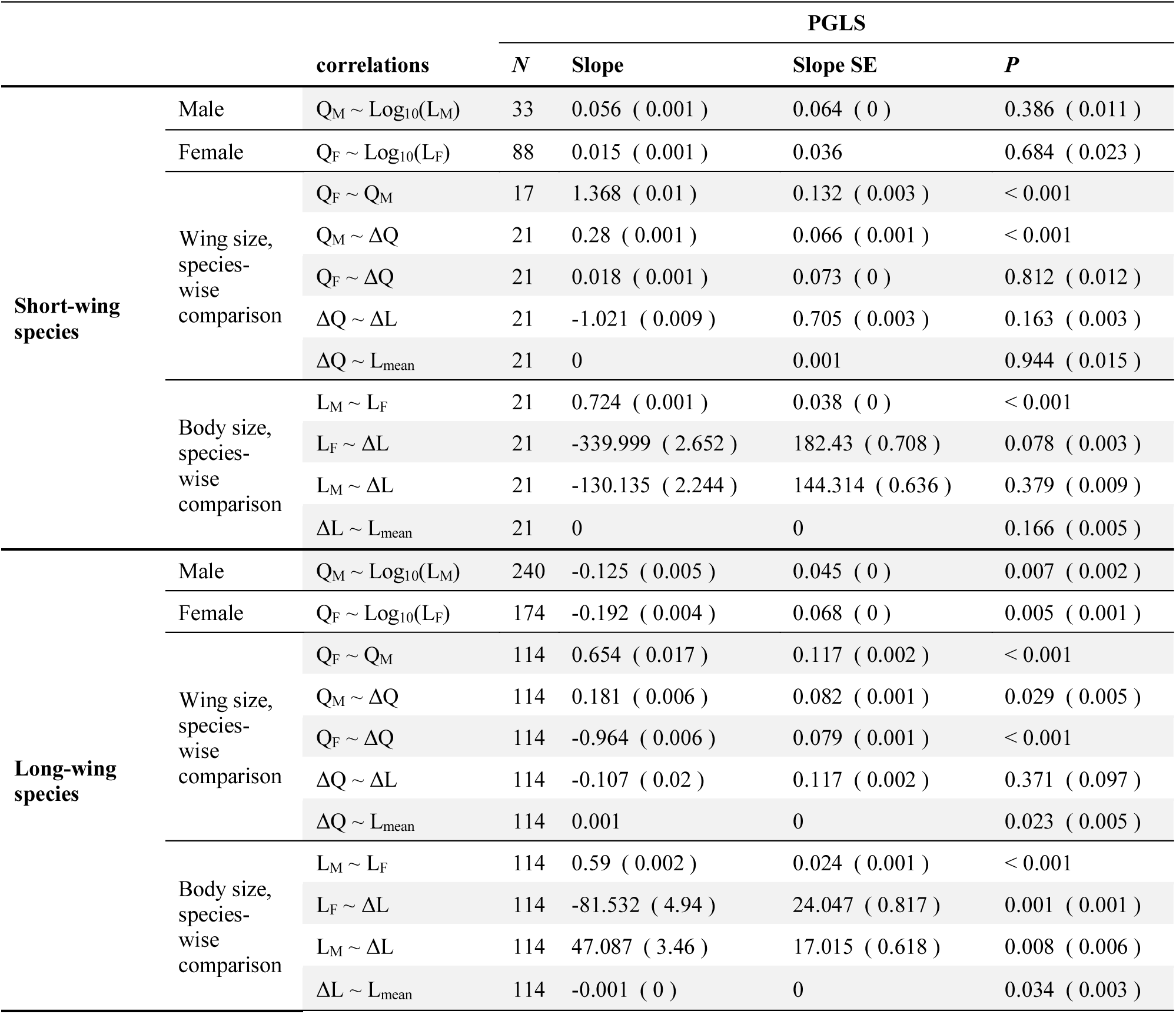
Summary of pairwise correlational analyses using PGLS. Values represent means from analyses using 100 randomly resolved trees, with 1 s.d. in brackets.

**Figure 7.**
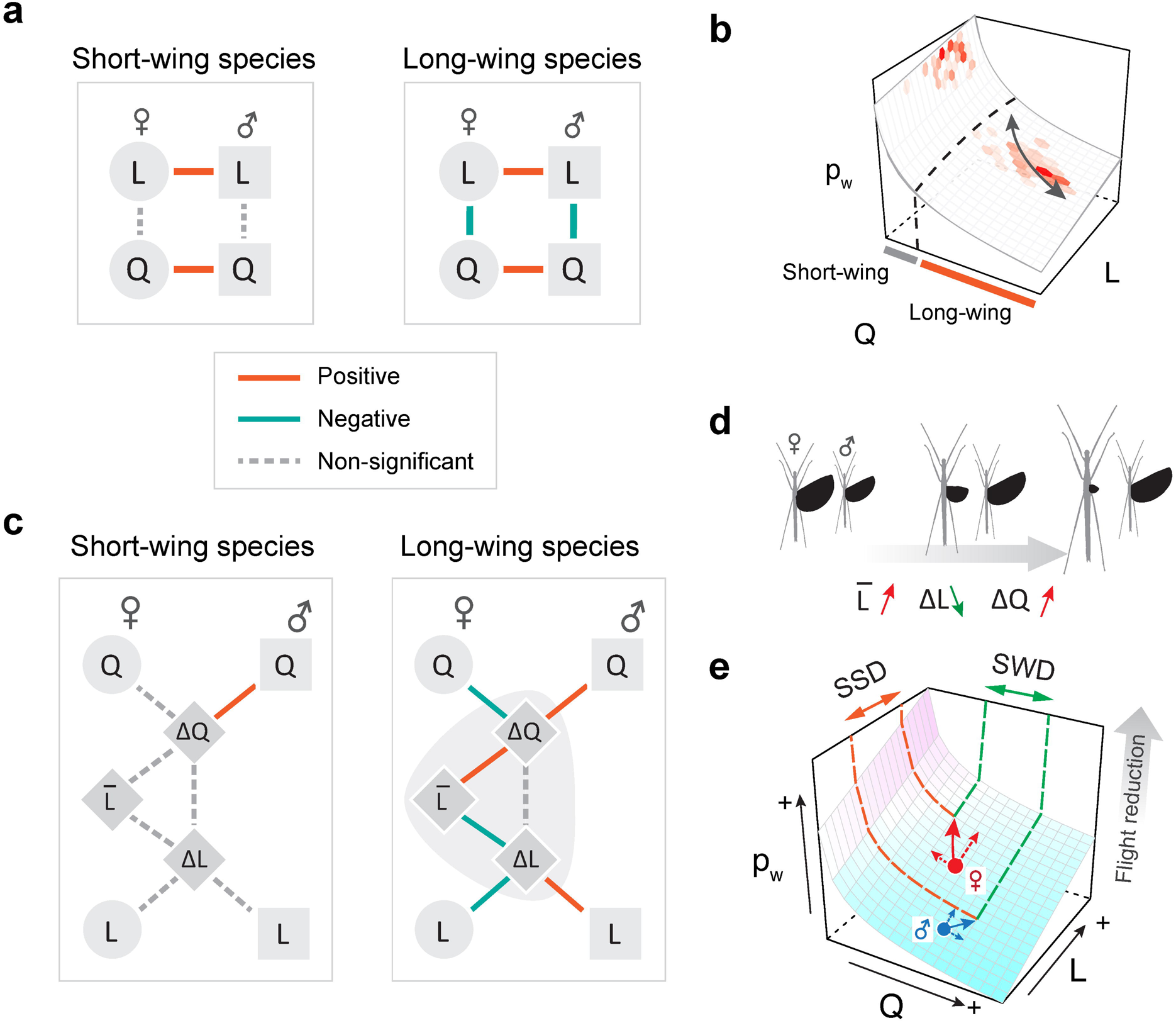
Phylogenetic correlations between wing and body size among stick insect species. (a) An inverse correlation between wing and body size was found in both sexes of long-winged species (Q > 0.3 in both sexes), indicated graphically in (b) as a coupled transition between different states of wing and body size on the wing loading landscape. (c) In long-winged species, sex-averaged L is significantly correlated with ΔL and with ΔQ, demonstrated graphically in (d) as an increasing sex-averaged L associated with decreasing SSD and increasing SWD. (e) Schematic scenario for consequences of increased sexual dimorphism in flight-related morphology; increases in female-biased SSD and male-biased SWD lead to greater difference in flight performance (i.e., changes in the *z*-position on the wing loading landscape). Details of the PGLS results are provided in **Table 3**.

## 4. Discussion

Most winged phasmid species possess either small or large wings (**Fig. 3b**). Few species have intermediate-sized wings, suggesting the presence of a fitness valley defined by two ‘adaptive peaks’ (see Stroud and Losos, 2016): one peak consists of wingless taxa and those with miniaturized wings (i.e., Q < ∼0.3), and another represents volant taxa (i.e., Q > ∼0.6). Insects with relative wing size near Q = ∼0.3 may be in transition between these two forms, with greater probability of either gaining or losing wing size depending on the interplay between various selective forces (see below). The predominance of wingless phasmid species may, in part, derive from reduced dispersal capacity leading to population isolation and ultimately genetic divergence. Given the possibility that repeated gain and losses of flight are associated with species diversification (Goldberg and Igić, 2008), linkage of evolutionary transitions between winged and wingless forms with diversification rates and overall macroevolutionary patterns should be addressed in future comparative studies. Significant wing size reduction over a relatively short divergence time, as in *A. tanarata* group, further demonstrates that the evolution of flightlessness can be recurrent and occur within nominal species. Similar scenarios of wing size reduction have been reported in alpine stoneflies (McCulloch et al., 2016). The evolution of flight-related morphology in phasmids can, in part, be viewed as displacement on the wing loading landscape (**Fig. 8a**), reflecting dual effects of variation in wing and body size. This multidimensional view provides a more complete perspective than consideration of wing size alone (as otherwise indicated by the inset arrows of **Fig. 8a**).

**Figure 8.**
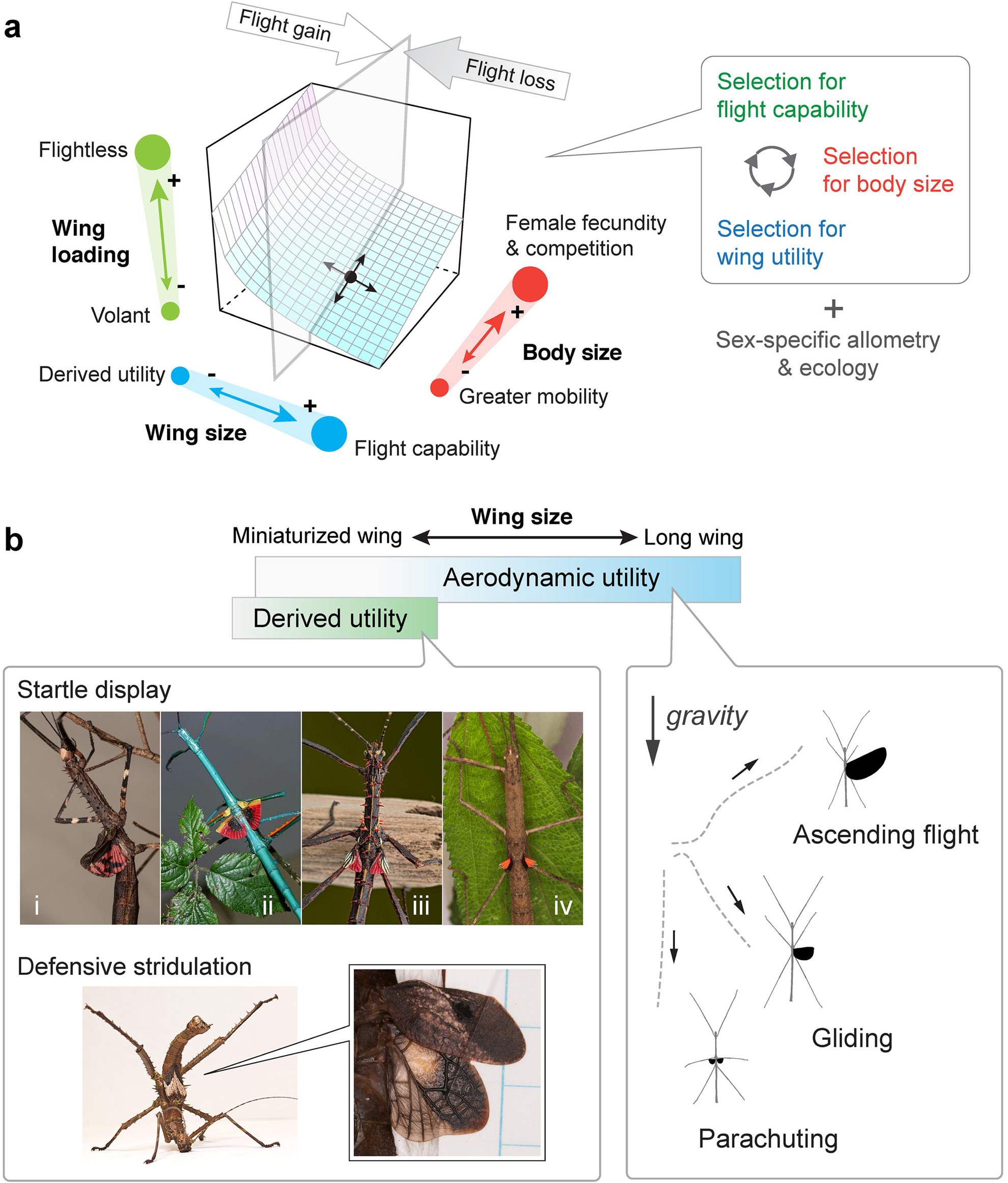
A summary of evolution of flight-related morphology in stick insects. (a) Schematic demonstration showing that the evolution of flight morphology (for any given position on the wing loading landscape) is driven by the interplay between three major forces and tradeoffs (inset). (b) Variation of wing utility with respect to relative wing size. Continuous variation in aerodynamic performance is coupled with the full spectrum of relative wing size variation, whereas derived functions such as use in startle displays or stridulation are frequently found in miniaturized wings. Examples of startle display: (i), *Diesbachia hellotis* female; (ii), *Achrioptera manga* male; (iii), *Parectatosoma* cf. *hystrix* male; (iv) *Oxyartes dorsalis* female (Photos of (i) – (iii) courtesy of Bruno Kneubühler). Example of defensive stridulation: *Haaniella echinata* male.

Flight in general enhances resource acquisition, dispersal, and escape from predators (Dudley, 2000), but wings can readily be lost in evolutionary time, or co-opted for non-aerodynamic purposes. Wing reduction in insects often derives from trade-offs with fecundity in particular contexts (e.g., habitat persistence, colonization of high-altitude environments; see Roff, 1994), whereas miniaturized and aerodynamically irrelevant wings often associate with derived defensive functions (e.g., startle displays and stridulation; Robinson, 1968; **Fig. 8b; Supplementary Table S3**). Altitudinal changes in life history strategies likely contribute to both body size miniaturization and wing reduction, as in the *A. tanarata* clade (**Fig. 2b**). For high-altitude species more generally, lower plant canopies at high elevations may reduce the functional significance of flight. By contrast, phasmid species with high dietary specificity might experience stronger selection for flight performance (e.g., Blüthgen et al., 2006). No data are presently available on flight abilities and associated aerodynamics among phasmid species.

For long-winged stick insects, aerial mobility may be an important component in sexual selection for enhanced male locomotor performance. Female phasmids tend to be less mobile and inconspicuous, whereas greater mobility in males may allow for greater success in dispersal and mating. The inverse correlation between wing and body size in male stick insects (**Fig. 7a,b**) suggests that selection for flight has limited the evolution of larger body size. Similar selection on male mobility and an enhanced locomotor apparatus has been documented in other male insects (Kelly et al., 2008). In wingless and short-winged species, larger body size might make a species more competitive in male-male competition (see Sivinski, 1978). A developmental tradeoff may limit the evolution of wing size, as shown by the inverse correlation between mating success and flight capability (as in Orthopteran and Hemipteran insects; Fujisaki, 1992; Crnokrak and Roff, 1995; Fairbairn and Preziosi, 1996). Future studies may compare variation in male body size between winged and wingless clades to test whether the evolution of male body size is constrained by selection for flight performance.

Sexual differences in mass allometry and body size are key factors influencing the evolution of phasmid wing dimorphism. Selection for increased fecundity will favor wing reduction in females, which can then lead to male-biased SWD as well as the evolution of defensive mechanisms that do not rely on flight; strong selection for flight capability may lead to female-biased SWD. Large female wings may be specifically favored in winged species with aerial copulation (e.g., *Trachythorax* spp.). Female-biased SSD is likely a canalized feature in orthopteroid insects, more generally (see Bidau et. al., 2016). In winged stick insects, the extent of SSD is clearly influenced by selection on fecundity in females, and by selection on flight in males. The allometric variation in SSD (i.e., the inverse correlation between ΔL and sex-averaged L; **see Fig. 7c,d**) is consistent with Rensch’s Rule (i.e., females are disproportionally larger in large species; Abouheif and Fairbairn, 1997; Fairbairn, 1997; Teder and Tammaru, 2005), instead of the converse outcome (i.e., isometric scaling in both sexes). This result may, however, be biased by allometric changes in body shape. For example, many phasmid species exhibit disproportionately slender bodies that may mimic plant stems, whereas other species have evolved thickened bodies for defense (e.g., the ‘tree lobster’ ecomorph; Buckley et al., 2008). In the scaling of wing loading (**Fig. 4d**), the contrast between *H. dilatata* male (family Heteropterygidae) and other phasmids (mostly in the subfamily Necrosiinae) suggests clade-specific allometry scaling. Future comparative assessment of body segment shapes and masses, in addition to body length, would enhance our understanding of allometric variation in SSD among phasmid taxa.

In winged phasmids, SSD and SWD are significantly correlated; this relationship does not pertain for either short- or long-winged species (**Fig. 7c**; **Supplementary Fig. S3**), reflecting interaction between multiple selective forces within sex-specific ecological contexts (**Fig. 8a**). The evolutionary intercorrelation between SSD and SWD is generally underexplored for most insects. Pterygote insects in general exhibit various types of SWD (e.g., male-biased and female-biased SWD have been reported in at least 11 and 5 orders, respectively; see Thayer, 1992), which can be correlated with sex-specific flight ecology (e.g., flight height and behavior; see DeVries et al., 2010) and with sexual selection for flight capability (e.g., copulation flight in caddisfly; Gullefors and Petersson, 1993). Future studies may address clade-specific SWD by correlating aforementioned factors within phylogenetic contexts.

These results for stick insects may provide more general insight into evolutionary transitions between wingless and fully winged insects. Given the widespread secondary loss of flight in pterygotes, sex-specific morphological scaling along the wing loading landscape can indicate the possible utility of partially reduced wings. Aerodynamic use of reduced wings during descent may be expected in arboreal pterygotes undergoing wing reduction, whereas non-aerodynamic functions would be predicted to be more likely in non-arboreal taxa (e.g., stridulatory wings in ground-dwelling insects). The eventual loss of aerodynamic utility may be characterized by a threshold wing loading (i.e., between 1 Nm^-2^ < p_w_ < 10 Nm^-2^ shown here), beyond which point selection for aerodynamic utility become insignificant. Similarly, morphological evolution associated with the origin of wings and of insect flight may have been sexually dimorphic, particularly if the earliest winglets served a non-aerodynamic function such as visual display (Alexander and Brown, 1963), with subsequently increases in size and mobility for aerial behaviors (see Dudley, 2000; Dudley and Yanoviak, 2011). Reductions in body size (with concomitantly lower wing loadings) may also favor the evolution of flight, as characterized the lineage leading to birds (Lee et al., 2014; Xu et al., 2014). Allometric variation in body structures can occur on both developmental and macroevolutionary timescales, and likely interacts with selection on aerodynamic performance. For example, if ancestral pterygotes retained winglets across nymphal instars, then selection for lower wing loading would foster allometric increases in wing size as well as a reduction in mass allometry (with less influence of body size increase to wing loading; **Fig. 5**). Physical models with wings of different sizes can be used to test biomechanical consequences of such differential allometries, as constrained by relevant morphologies inferred from the fossil record. And for extant phasmids, assessment of flight behaviors and aerial capacity across taxa is now clearly warranted.

## Author contributions

Y.Z. devised experiments and performed the majority of data collection and analyses. C.O. and S.P. participated in morphological data collection. S. S. contributed to molecular data collection. X. C. contributed to phylogenetic analyses. F. H. contributed to field data collection of *Asceles tanarata* group. All authors contributed to writing of the manuscript.

## Acknowledgments

We thank Francis Seow-Choen for providing captive bred insects, helping with field works, and commenting on stick insect natural history. We also thank Lynn Nguyen, Camille Gonzales, Faye Pon, Joan Chen, Nina Gnong, Stephanie Yom, Yuexiang Chen, Chulabush Khatancharoen, Biying Li, Grisanu Naing, Jizel Emralino, Xiaolin Chen, Ian Abercrombie, Yamai Shi-Fu Huang, Azuan Aziz and Juhaida Harun for help with data collection, and David Wake and Sven Bradler for comments on the manuscript. We further thank Paul Brock, Nicholas Matzke and Emma Goldberg for comments at the early stage of the work, and the Forestry Department of Pahang, Malaysia, for permission of insect collection. This research was supported by National Science Foundation (DDIG 1110855), Museum of Vertebrate Zoology, the Department of Integrative Biology at UC-Berkeley, the Undergraduate Research Apprentice Program (URAP) of UC-Berkeley, the Society for Integrative and Comparative Biology (SICB), and the Ministry of Higher Education Malaysia (FRGS/1/2012/SG03/UKM/03/1(STWN)).

## Symbols and abbreviations

A_w_: Wing area
p_w_: Wing loading
L: Body length
L_w_: Wing length
m: Mass
SSD: Sexual size dimorphism
SWD: Sexual wing dimorphism
Q: Relative wing size
ΔL: Sexual size dimorphism index
ΔQ: Sexual wing dimorphism index

