## Supplementary Information for "A tale of winglets: evolution of flight morphology in stick insects"

#### Supplementary Materials

**Text section**

Notes on ancestral state reconstruction for relative wing size in stick insects

**Figure**

S1, Summary of morphological sampling
S2, Polytomous tree with mapped traits
S3, Ancestral state reconstruction for relative wing size
S4, Ancestral state reconstruction for body size
S4, Ancestral state reconstruction for sexual size dimorphism (SSD) and sexual wing dimorphism (SWD)
S5, Phylogenetic mapping of hindwing state for sampled tribes
S6, Summary of phylogenetic correlations for flight morphology among winged stick insects

**Table**

S1, Information on nuclear and mitochondrial loci
S2, Summary of PGLS results for all winged stick insects
S3, Summary of derived wing functions for stick insects with miniaturized wings

**Datasets 1-4**

1. Morphometrics (taxon, wing length, body length)
2. Scaling of wing loading (body mass, wing length, wing area)
3. Molecular sampling (list of species), including accession numbers for new sequences on GenBank
4. Nexus input files

**Script**

MatLab script for organizing taxonomic data from Phasmida Species Files

### Supplementary Text

#### Notes on ancestral state reconstruction for wing size evolution in stick insects

Our maximum-likelihood ancestral state reconstruction showed that a short relative wing size ($Q < 0.4$) preceded multiple gains and losses observed in extant phasmids (**Fig. S4,5**). Nevertheless, this result offered only a preliminary perspective on wing size evolution in phasmids, as it reflects but a small portion of extant phasmid diversity (~ 10% documented species). A complete molecular phylogeny and comprehensive trait sampling are not currently available.

Preliminary sampling for the presence of hindwings showed a highly variable pattern across phasmid clades, even below the tribe level (**Fig. S7**). In order to properly model evolutionary rates of change and to reconstruct ancestral states for relative wing size, future work should increase phylogenetic resolution and incorporate trait sampling across extant tips. Properly modeling gain and loss of wings in phasmids will require complex models to account for (1) variation in the rate of wing size evolution across the tree (Beaulieu et al. 2013), and (2) variation in rates of species diversification across the tree (Maddison 2006). In particular, Goldberg and Igić (2008) suggested that the loss and gain of wings may affect diversification patterns. Our results demonstrating a bimodal distribution of wing size further indicated a more complex and wing size-dependent diversification scenario.

#### Supplementary Figures

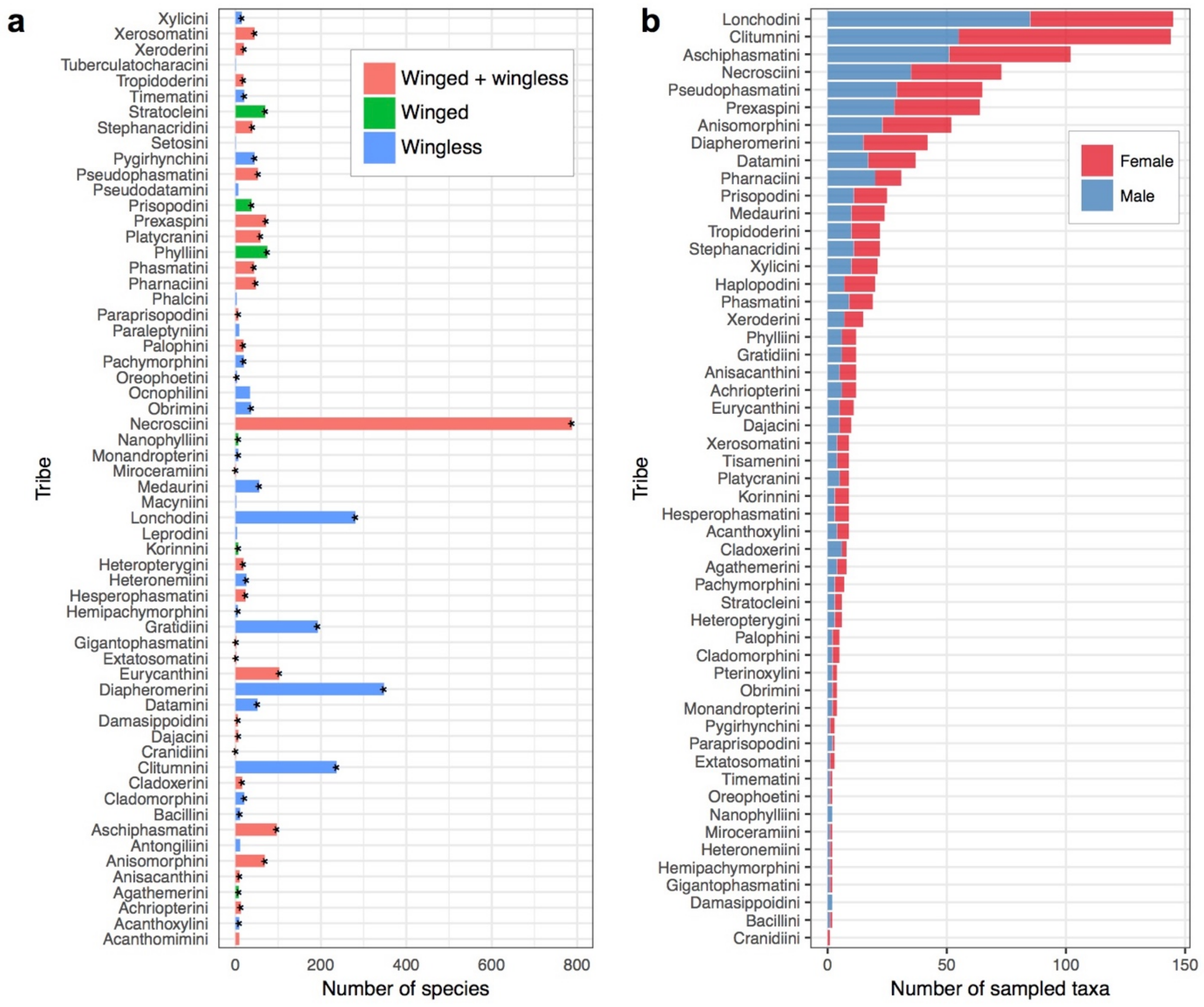


**Figure S1. Summary of taxa sampled for flight-related morphology.** (a) All documented phasmid tribes, with the number of species based on Brock et al. (2019). Colors represent the hindwing condition within tribes; asterisks denote tribes sampled in this study. (b) Summary of the number of taxa sampled for hindwing size (see **Supplementary Dataset 1**).

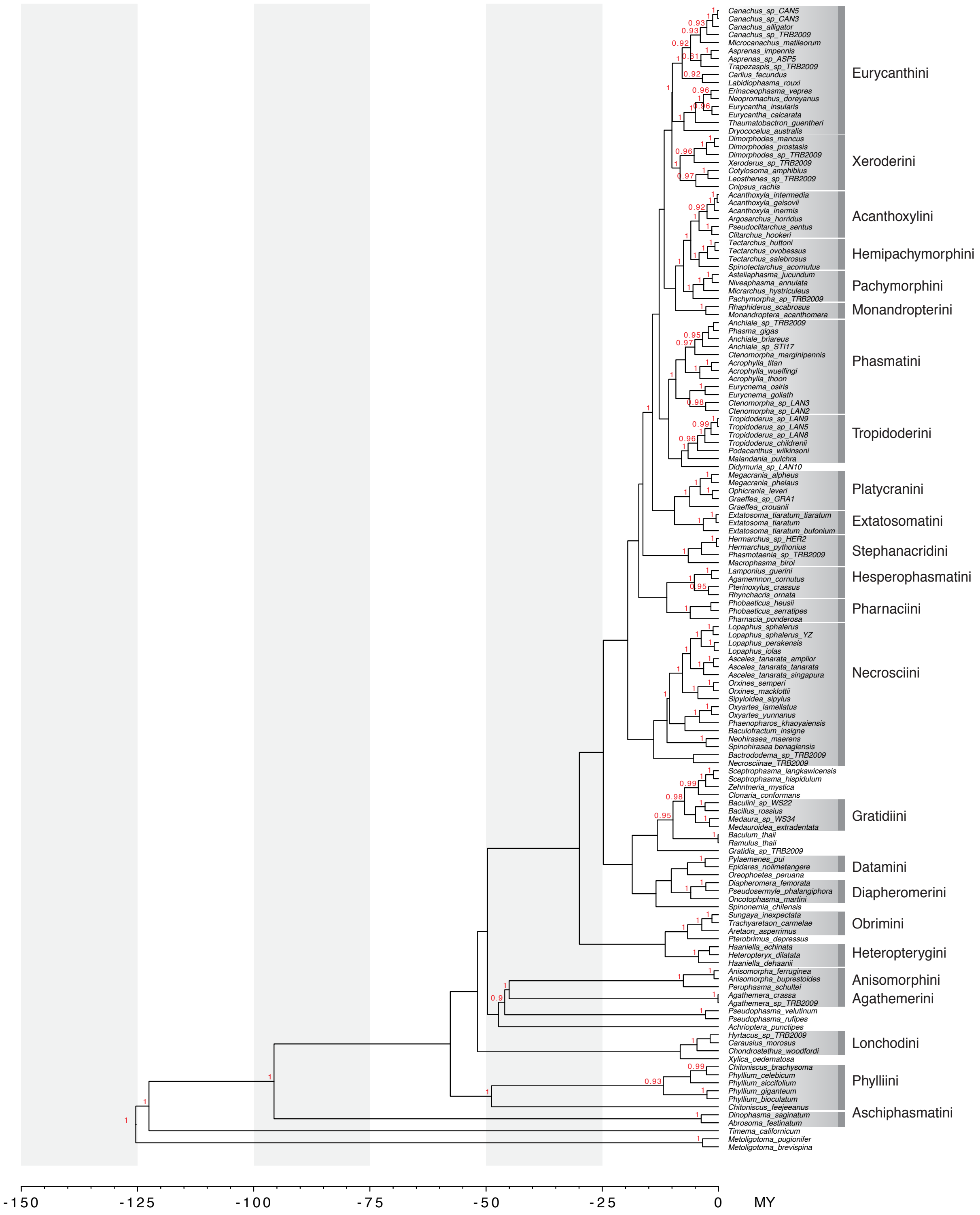


**Figure S2.** Bayesian species tree inferred with BEAST, with node labels showing posterior probabilities > 0.9.

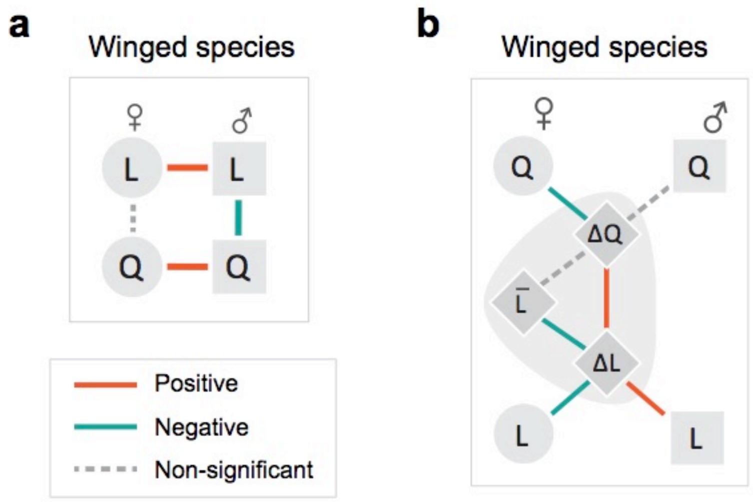


**Figure S3.** Phylogenetic correlations between wing and body size among winged species. Details are provided in **Table S2**.

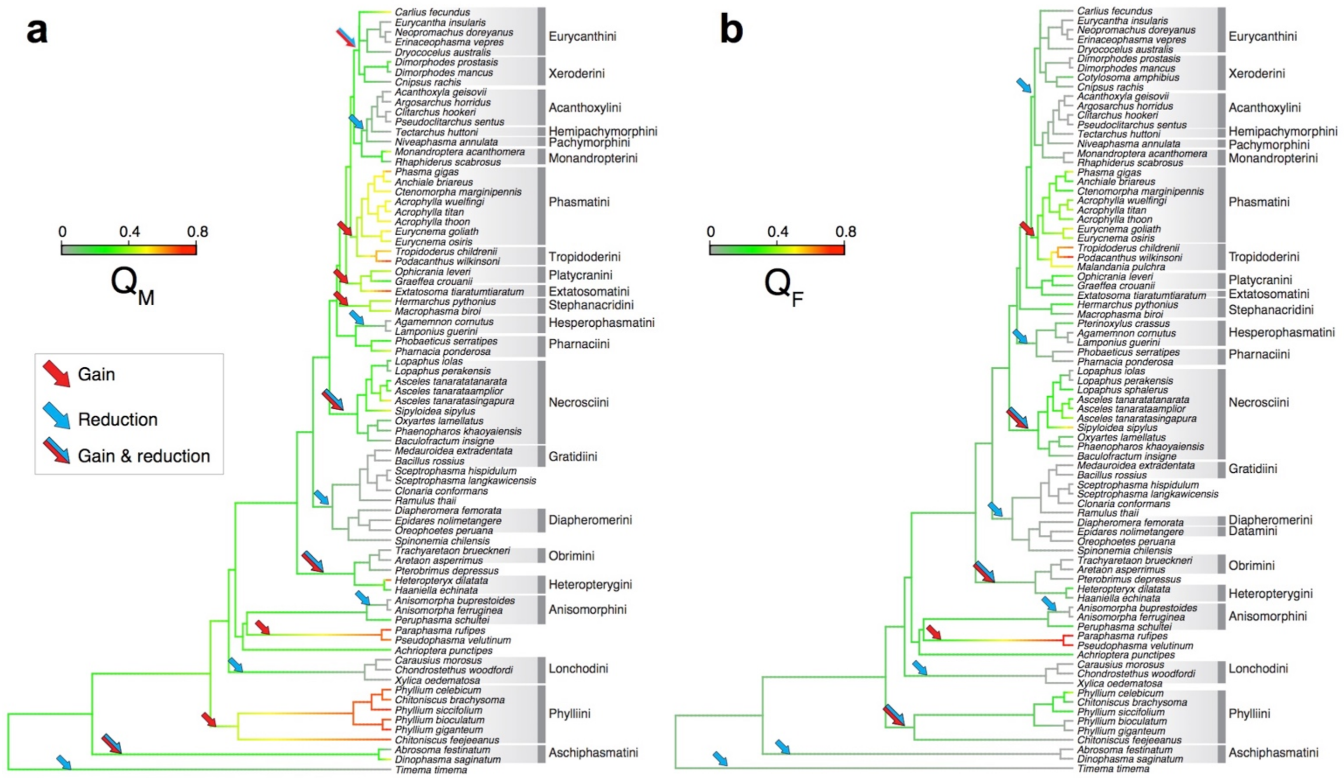


**Figure S4**. Ancestral state reconstruction for relative wing size (Q) of males (a) and females (b), respectively, featuring repeated gain and reduction. The last common ancestor of the main clades is characterized by short wings (Q < 0.4); however, this inference is liable to change with more complete sampling. Arrows on main branches highlight increases and reductions of wing size. In long-wing lineages (e.g., Tropidoderini and Necrosciini), relative wing size between females and males is correlated (see **Fig. 7a**).

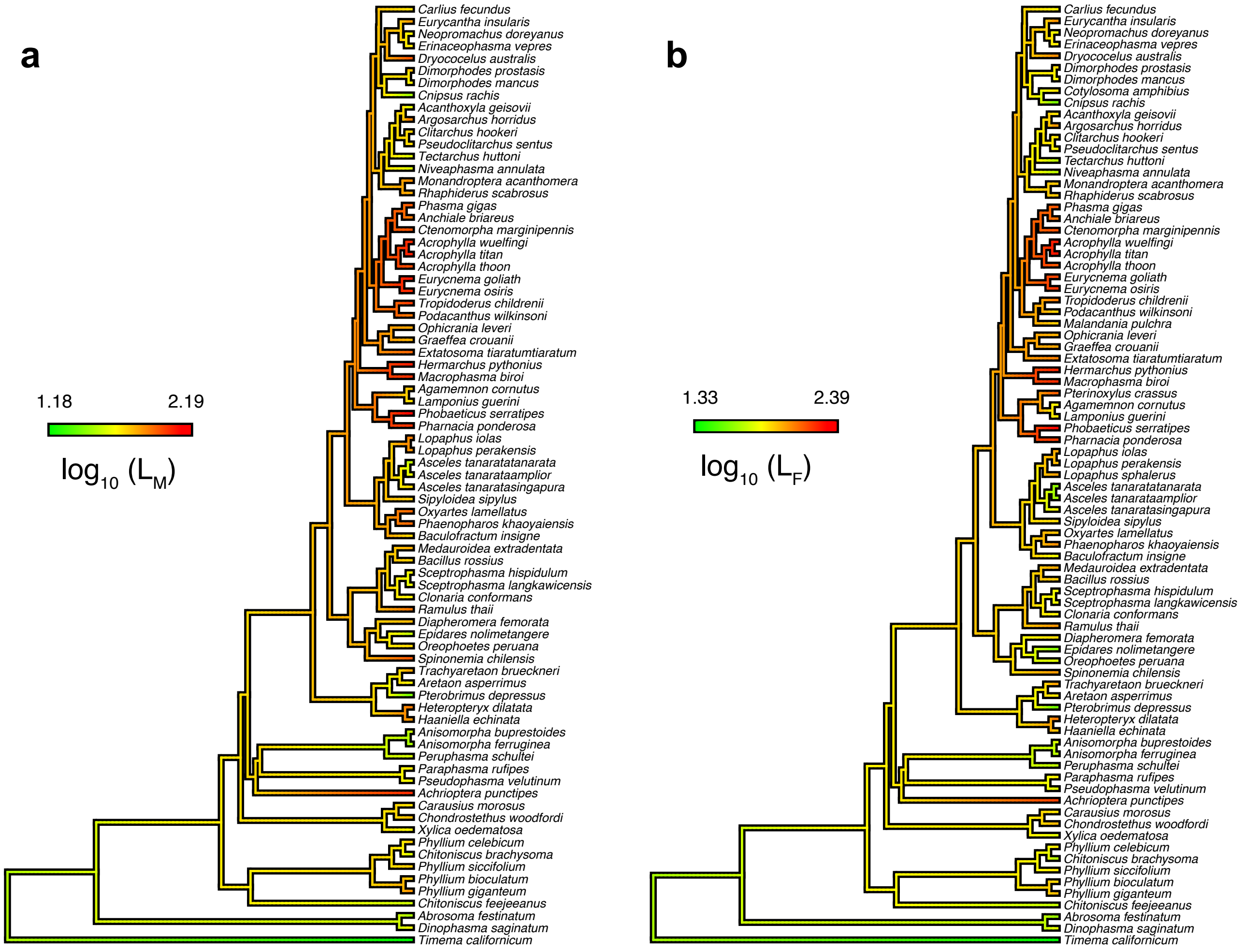


**Figure S5**. Ancestral state reconstruction for body size (L) of males (a) and females (b), respectively. Body size evolution is coupled between two sexes but independent of wing size evolution, corresponding with the correlational pattern summarized in **Fig. 7a**.

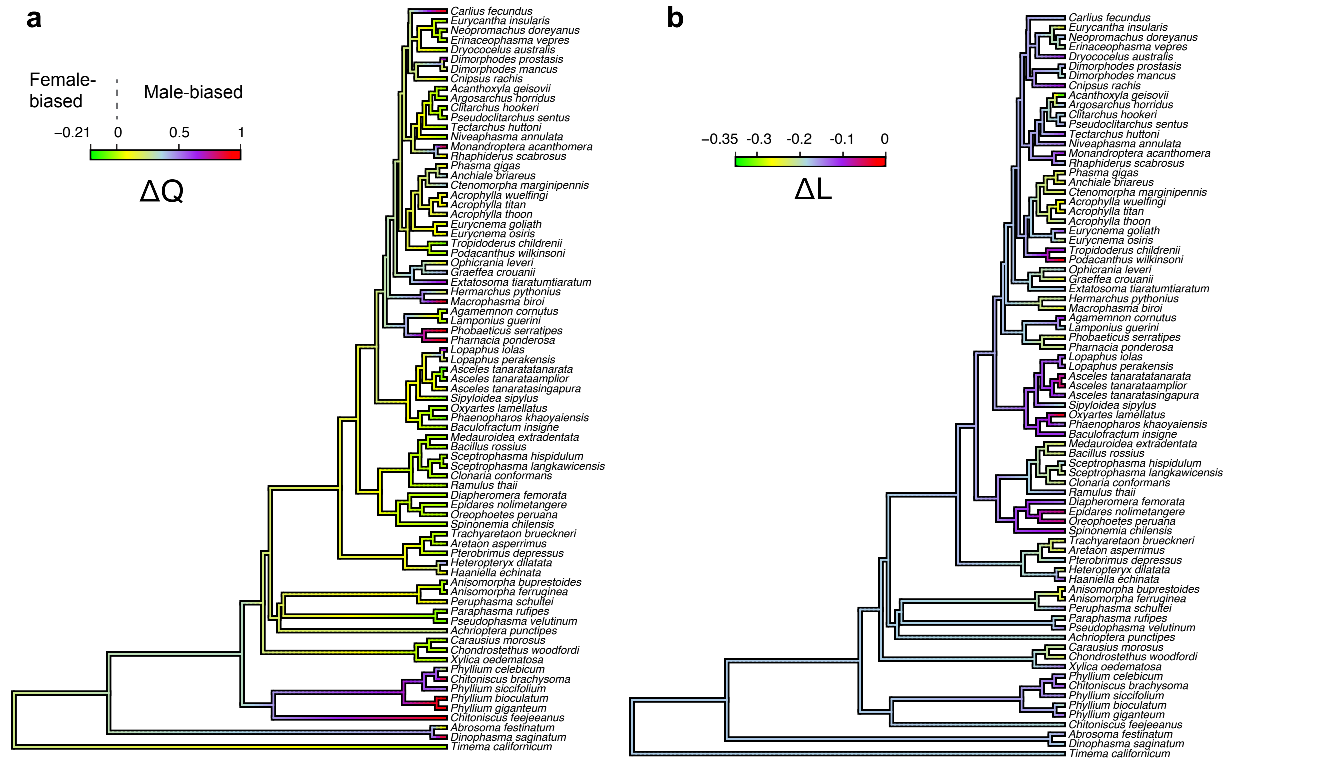


**Figure S6**. Ancestral state reconstruction for sexual wing dimorphism (SWD) index (ΔQ; a) and sexual size dimorphism (SSD) index (ΔL; b). (a) The ancestral state reconstruction of ΔQ shows repeated increases and reductions, with an intermediate level of male-biased SWD ($0 < \Delta Q < 0.5$, as represented by yellow color) as the ancestral state for most clades. (b) Ancestral state reconstruction of ΔL, showing an intermediate level of female-biased SSD ($-0.2 < \Delta L < -0.1$) as the ancestral state preceding repeated increase and reduction. The lack of evolutionary correlation between ΔL and ΔQ, as summarized in **Fig. 7c**, is evident by comparing (a) and (b).

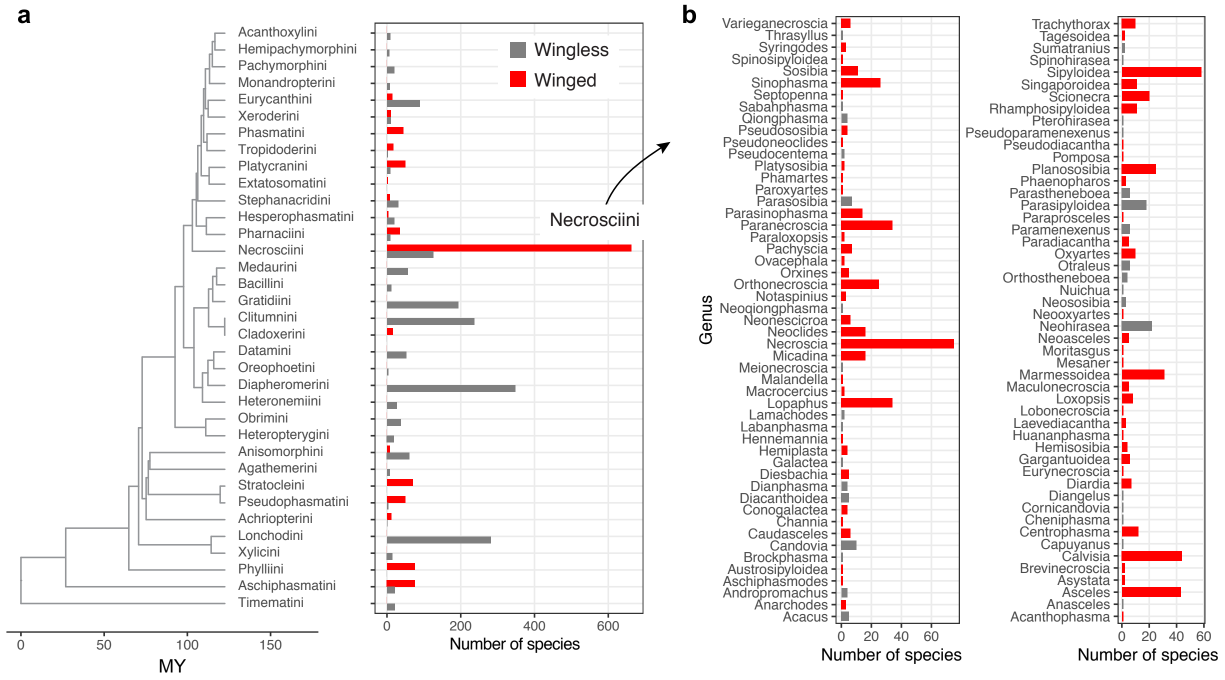


**Figure S7**. Summary of species richness for tribes represented by the molecular phylogeny. (a) The number of known species mapped on a phylogeny of tribes sampled in this study, with colored bars showing the number of species with and without hind wings. (b) A more detailed summary of the number of known species and the presence of wings for Necrosciini, the most diversified tribe. Taxonomic organization followed Phasmida Species File (Brock, 2019).

### Supplementary Table

| Locus | Forward primer | Reverse primer | $T_a$ (˚C) | Length (bp) | Reference |
|---|---|---|---|---|---|
| 28s | AGAACTTTGAAGAGAGAGTTCAAGA | TCAAGACGGGTCGGGAGA | 50 | 495 | Buckley et al., 2008 |
| COI | TTGATTTTTTGGTCATCCAGAAGT | TCCAATGCACTAATCTGCCATATTA | 50 | 814 | Simon et al., 1994 |
| COII | AATATGCAGATTAGTGCA | GTTTAAGAGACCAGTACTTG | 50 | 736 | Simon et al., 1994 |
| H3 | ATGGCTCGTACCAAGCAGAC | ATATCCTTRGGCATRATRGTGAC | 50 | 358 | Buckley et al., 2008 |

**Table S1.** Information on the nuclear and mitochondrial loci used in this study.

| | | PGLS | | | |
|---|---|---|---|---|---|
| | correlations | *N* | slope | slope.SE | *P* |
| **Male** | $Q_M$ ~ $Log_{10}(L_M)$ ‡ | 273 | -0.271 ( 0.005 ) | 0.072 ( 0.001 ) | < 0.001 |
| **Female** | $Q_F$ ~ $Log_{10}(L_F)$ ‡ | 262 | -0.107 ( 0.02 ) | 0.1 ( 0.001 ) | 0.296 ( 0.096 ) |
| **Wing size, species-wise comparison** | $Q_F$ ~ $Q_M$ ‡ | 158 | 0.876 ( 0.01 ) | 0.063 ( 0.001 ) | < 0.001 |
| | $Q_M$ ~ ΔQ | 183 | -0.029 ( 0.003 ) | 0.038 ( 0 ) | 0.438 ( 0.04 ) |
| | $Q_F$ ~ ΔQ ‡ | 158 | -0.534 ( 0.007 ) | 0.068 ( 0 ) | < 0.001 |
| | ΔQ ~ ΔL ‡ | 158 | -0.637 ( 0.033 ) | 0.22 ( 0.003 ) | 0.005 ( 0.003 ) |
| | ΔQ ~ $L_{mean}$ ‡ | 158 | 0.001 ( 0 ) | 0.001 | 0.332 ( 0.11 ) |
| **Body size, species-wise comparison** | $L_M$ ~ $L_F$ | 367 | 0.61 ( 0.001 ) | 0.014 ( 0 ) | < 0.001 |
| | $L_F$ ~ ΔL | 367 | -118.893 ( 5.122 ) | 18.92 ( 0.38 ) | < 0.001 |
| | $L_M$ ~ ΔL | 367 | 31.48 ( 3.534 ) | 13.999 ( 0.276 ) | 0.03 ( 0.02 ) |
| | ΔL ~ $L_{mean}$ | 158 | 0 | 0 | 0.03 ( 0.026 ) |

**Table S2**. Summary of pairwise correlational analyses applied to all winged species using PGLS models. Values represent means from analyses using 100 randomly resolved trees, with 1 s.d. in brackets. The symbol ‡ indicates that equivalent significance was found for analyses using the original data with variables converted to pseudo-continuous ordinal numbers in analyses (see Methods).

| Taxa | Stridulation | Startle display | Reference |
|---|---|---|---|
| *Pterinoxylus spinulosus* female and male | x | x | Robinson, 1968 |
| *Hanniella spp.* female and male | x | | Pers. Observ. |
| *Oxyartes spp.* female and male | | x | Pers. Observ. |
| Phaenopharus spp. female and male | | x | Pers. Observ. |
| *Diapherodes spp.* female | | x | Pers. Observ. |
| *Parectatosoma spp.* female and male | | x | Pers. Observ. |
| *Diesbachia hellotis* female | | x | Pers. Observ. |
| *Achrioptera spp.* female and male | | x | Pers. Observ. |
| Peruphasma spp. female and male | | x | Pers. Observ. |

**Table S3.** A summary of known cases of derived wing utility in stick insects with miniaturized wings.
